# Design of a stem cell-based therapy for ependymal repair in hydrocephalus associated with germinal matrix hemorrhages

**DOI:** 10.1101/2023.04.13.536749

**Authors:** Luis M. Rodríguez-Pérez, Betsaida Ojeda-Pérez, María García-Bonilla, Javier López-de San Sebastián, Marcos González-García, Beatriz Fernández-Muñoz, Rosario Sánchez-Pernaute, María L. García-Martín, Dolores Domínguez-Pinos, Casimiro Cárdenas-García, Antonio J. Jiménez, Patricia Páez-González

## Abstract

Germinal matrix hemorrhages (GMH) and the consequent posthemorrhagic hydrocephalus (PHH) are among the most common and severe neurological complications of preterm birth that require lifelong complex neurosurgical care. GMH and PHH provoke disruption of neuroepithelium/ependyma development, a key structure implicated in brain development and homeostasis. Neuroepithelial/ependymal damage causes lifelong cognitive and motor deficits; however, no therapy is directed to recover the damaged ependyma. This study is aimed to test the possibilities of ependymal repair in GMH/PHH using neural stem cells (NSCs) or ependymal progenitors (EpPs). Thus, it sets the basis for a therapeutic approach to treating ependymal damage and preventing brain developmental deficits. GMH/PHH was induced in 4-day-old mice using different experimental procedures involving collagenase, blood, or blood serum injections. PHH severity was characterized using magnetic resonance, immunofluorescence, and protein expression quantification with mass spectrometry. Additionally, a new *exvivo* approach using ventricular walls from mice developing moderate and severe GMH/PHH was generated to study ependymal restoration and wall regeneration after stem cell treatments. NSCs or EpPs obtained from newborn mice were transplanted in the explants, and pretreatment with mesenchymal stem cells (MSCs) was tested. Ependymal differentiation and the effect of MSC-conditioned microenvironment were investigated in both explants and primary cultures. In the animals, PHH severity was correlated with the extension of GMH, ependymal disruption, astroglial/microglial reactions, and ventriculomegaly. In the explants, the severity and extension of GMH hindered the survival rates of the transplanted NSCs/EpPs. In the explants affected with GMH, new multiciliated ependymal cells could be generated from transplanted NSCs and, more efficiently, from EpPs. Blood and TNFα negatively affected ciliogenesis in cells expressing Foxj1. Pretreatment with mesenchymal stem cells (MSC) improved the survival rates of EpPs and ependymal differentiation while reducing the edematous and inflammatory conditions in the explants. In conclusion, in GMH/PHH, the ependyma can be restored from either NSC or EpP transplantation, being EpPs in an MSC-conditioned microenvironment more efficient for this purpose. Modifying the neuroinflammatory microenvironment by MSC pretreatment positively influenced the success of the ependymal restoration.

## Introduction

Germinal matrix hemorrhages (GMH) and subsequent intraventricular hemorrhages (IVH) are major causes of morbidity and mortality in the premature neonatal population and require lifelong, complex neurosurgical care^1, 2^. One of the major complications of GHM/IVH is the development of posthemorrhagic hydrocephalus (PHH) with progressive enlargement of the head and the ventricular system^3, 4^. Thus, a large GMH/IVH is associated with an increased risk of adverse neurologic sequelae^3, 5^. Advances in neonatal medicine have led to a higher survival rate in preterm infants. However, the rate of GMH/IVH has also increased^6^. Furthermore, disabilities and mortality rates in PHH cases are higher than those in other forms of pediatric hydrocephalus^7^.

PHH injuries include periventricular edema and neuroinflammation, demyelination, axonal degeneration and impaired axoplasmic transport in periventricular white matter, cerebral hypoxia and ischemia, affected blood‒ brain and blood–cerebrospinal fluid (CSF) barriers, and the presence of altered levels of neurodevelopmental trophic factors^8–11^. In these conditions, hydrocephalus development is associated with reactions in astrocytes and microglia^12, 13^.

Disruption of multiciliated ependyma development^12, 14–16^ is a relevant and key event associated with GMH/IVH and PHH. The ependyma constitutes a cell barrier between the brain parenchyma and the ventricle CSF^17–19^. Ependymal disruption and gliosis in PHH can affect CSF circulation and reabsorption^1, 17^. Furthermore, ependymal loss can lead to obstructive hydrocephalus, as described in experimental hydrocephalus and human fetuses^20–22^. In the cerebral aqueduct, ependymal loss influences the disease outcome^20^. In addition, the ependyma is a key player in regulating neurogenesis^23, 24^. Iron and other blood components can be involved in ependymal disruption and dysfunction and, accordingly, in the development of PHH after GMH/IVH^25^. Lysophosphatidic acid in the blood serum appears to be implicated in the disruption of ependymal development^26, 27^. Peroxiredoxin 2, an essential protein in cell metabolism, is abundant in erythrocytes and disrupts ependymal development, leading to hydrocephalus^28^. In this way, ependyma can be considered a key therapeutic target because of its relevance for hydrocephalus occurrence and role in CSF circulation and physiology^17–19, 29–31^. Nevertheless, no therapy is aimed to functionally recover nor regenerate this main structure for brain development and homeostasis, leading to an essential clinical therapeutic deficit.

Pediatric hydrocephalus evolves through adulthood, and surgical treatment results in more complications in PHH and therefore in its outcome^1, 32, 33^. Treatments are mostly restricted to alleviating ventricular pressure by CSF drainage through ventricular shunts, third ventriculostomy, or extraventricular drains^33, 34^. Thus, these treatments are only palliative and can only prevent some PHH-associated damage. Treatments are also being sought to prevent the development of GMH, although the molecular causes of its development are not well understood^35, 36^. Attempts to develop nonsurgical therapies for pediatric hydrocephalus have not been successful^37^. Therefore, new approaches or tools are needed to provide neuroprotection and promote the repair or regeneration of damaged tissues^38^, especially the regeneration of the ependyma. Stem cell-based therapies appear promising for several neurodegenerative diseases due to their regenerative potential^39^. In this line, various attempts using stem cell-based therapies have been assayed in experimental forms of congenital and acquired fetal-neonatal hydrocephalus. In the case of neural stem cells (NSCs)^40, 41^ and mesenchymal stem cells (MSCs)^42–46^, promising results have been reported in hydrocephalus of obstructive and posthemorrhagic origins. However, in hydrocephalus after IVH, although the application of umbilical cord stem cells could attenuate PHH, the ependyma did not regenerate^42, 44, 47^.

This study is aimed to test the possibility of directly repairing the ependyma in PHH using NSCs and ependymal progenitor (EpPs) cells. For this purpose, PHH was modeled in neonatal mice at a developmental stage equivalent to that when neurologic processes are affected in human cases after GMH/IVH^48^. Different experimental approaches have been developed in animals to replicate the pathology in PHH using intracerebroventricular blood injection or GMH induction by collagenase injection. The latter has been proven to mimic human PHH^4, 49^. In the present investigation, blood serum intracerebroventricular injection was also tested given the effect of LPA on ependyma development and a possible role in PHH etiology. In this study, mice exhibited two different forms of PHH. A group of mice exhibited a severe form of PHH (sevPHH) with pronounced ventriculomegaly and severe damage to the white matter. In contrast, another group of animals presented mild ventricular dilatation considered moderate PHH (modPHH). The extension of ependymal disruption was correlated with PHH severity. We also found that both forms of PHH shared common cytopathology, including disruption of ependymal development, glial reactions, and myelin affectation. Additionally, a new *ex vivo* approach using ventricular wall explants of mice with moderate and severe GMH/PHH (modGMH/PHH and sevGMH/PHH) was generated to test stem cell therapy and the possibility of ependymal restoration. With this approach, it has been possible to investigate the differential therapeutic potential of EpPs and NSCs after transplantation in *ex vivo* conditions and considering the severity of the neuropathological and neuroinflammatory conditions. In addition, it has been tested how inflammatory microenvironment modification by bone marrow-derived MSC pretreatment improves the possibility of ependymal repair by NSCs and EpPs.

## Results

### Generation of moderate and severe posthemorrhagic hydrocephalus

Although there are different strategies to develop PHH experimentally, a systematic method able to give rise to PHH that mimics the human disease with different degrees of neuropathological damage and ependymal alteration was needed. Therefore, we decided to generate PHH/GMH by injection with whole blood, blood serum, and collagenase.

With the different methodological procedures, mice exhibiting severe PHH (sevPHH), which presented at least a fiftyfold increase in the lateral ventricle volume, were obtained 14 days after injection with blood serum into the right lateral ventricle (2.75% of 11 total mice) or both lateral ventricles (7.69%, of 11 total mice), whole blood into the right lateral ventricle (18.2% of 13 total mice), or collagenase into the germinal matrix in both hemispheres (55.5% of 33 total mice, 5 mice died a day after injection; therefore these animals could not be classified into any group) (Fig. 1a, b). In addition, a significant percentage of mice at the same age developed only mild ventriculomegaly in the lateral ventricles, with at least a twofold volume increase compared with that of normal mice, when injected with blood serum in the right lateral ventricle (54.5%) or both lateral ventricles with blood serum (61.5%), whole blood, (27.3%), and collagenase (30.3%) (Fig. 1a, b). These latter mice were considered to have a moderate form of PHH (modPHH), as confirmed by histopathological and protein expression analyses described below.

**Fig. 1.**
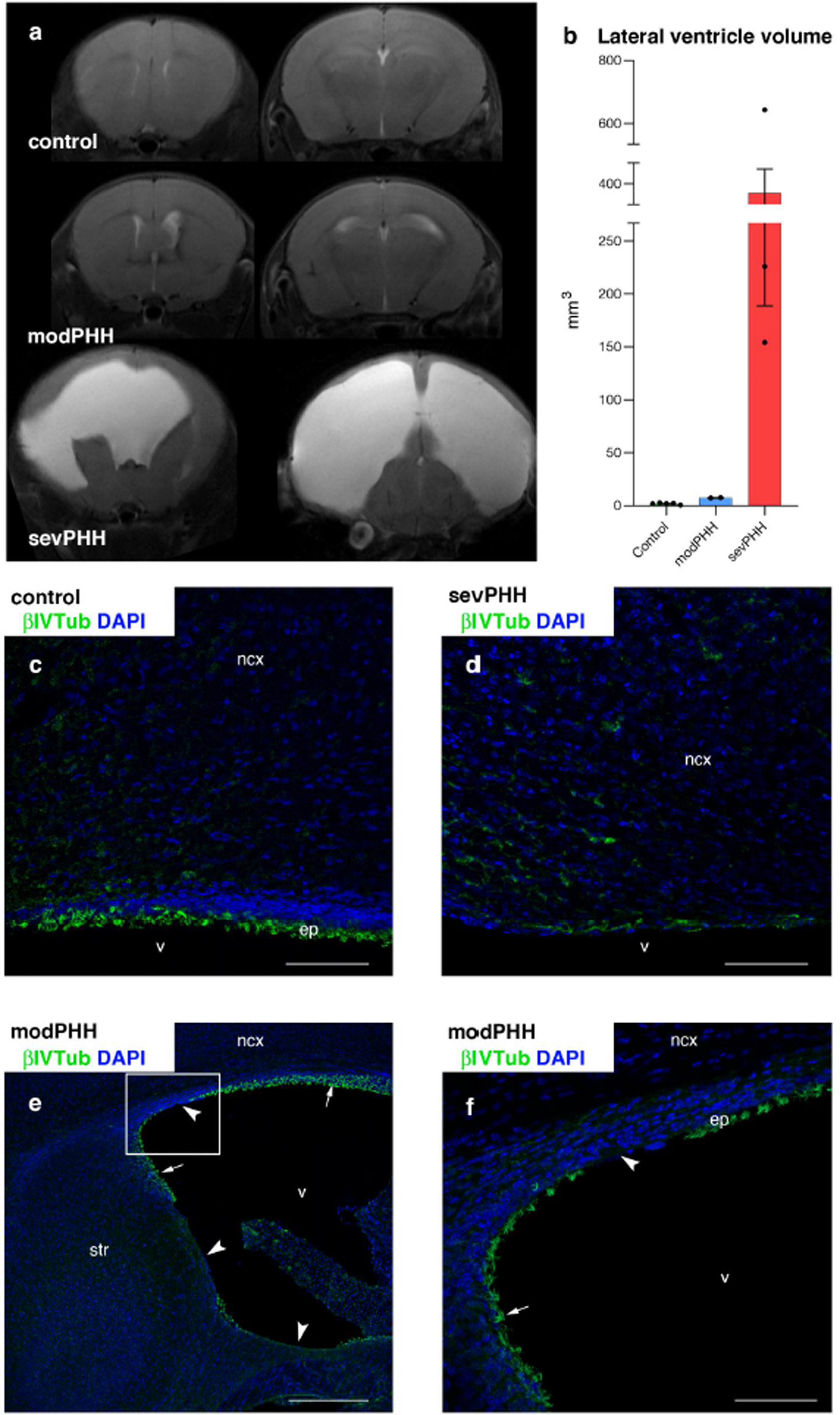
Differential lateral ventricle enlargement after GMH induction by collagenase injection. Mice exhibited mild ventriculomegaly (*modPHH*) and large ventriculomegaly (*sevPHH*) compared to a normal mouse (*control*). **a** T2 magnetic resonance images showing two different levels of representative mice with modPHH and sevPHH and a normal control mouse. **b** Representative examples of lateral ventricle volume for animals from the different experimental groups, mean ± SEM (control, n = 5; moPHH, n = 2; sevPHH, n = 3). **c-f** Ependymal denudation (*arrowheads*) in the lateral ventricle detected by cilia immunostaining with βIV-Tubulin (*βIVTub*, *arrows*). Cell nuclei are stained with DAPI (*blue*). Abbreviations: *str*, striatum. Scale bars: **c**, **d**, **f**, 77 µm; **e**, 300 µm.

### Mice with moderate and severe forms of posthemorrhagic hydrocephalus shared common neuropathological events

Once GMH/PHH was generated by injection with whole blood, blood serum, and collagenase, the effect on the ventricular walls was investigated. The same neuropathological events were detected after injection of blood serum into both lateral ventricles, whole blood into the right lateral ventricle, or collagenase into the two hemispheres in the germinal matrix. Because of the higher efficiency of collagenase injections to produce GMH/PHH and how it mimics human disease^49^, the neuropathology of this model was thoroughly analyzed and described below.

Mice with modPHH and sevPHH (Fig. 1c-f) showed different extensions of lateral ventricle surfaces denuded of ependyma (sevPHH, n = 5, mean = 88.22%, SD ± 10.09; modPHH, n = 4, mean = 19.04% of the lateral ventricle surface, SD ± 8.34; p < 0.0001, t = 11, df = 7, unpaired two-tailed Student’s t test). In addition, mice with both sevPHH and modPHH presented a periventricular white matter astrocyte reaction associated with ependyma-denuded surfaces (Fig. 2a-c). Moreover, microglial reactions were present overall in the ventricle wall of the mice with sevPHH. Still, in the case of modPHH, they appeared to be more evident in the ventricle walls affected by ependymal disruption (Fig. 2d-f).

**Fig. 2.**
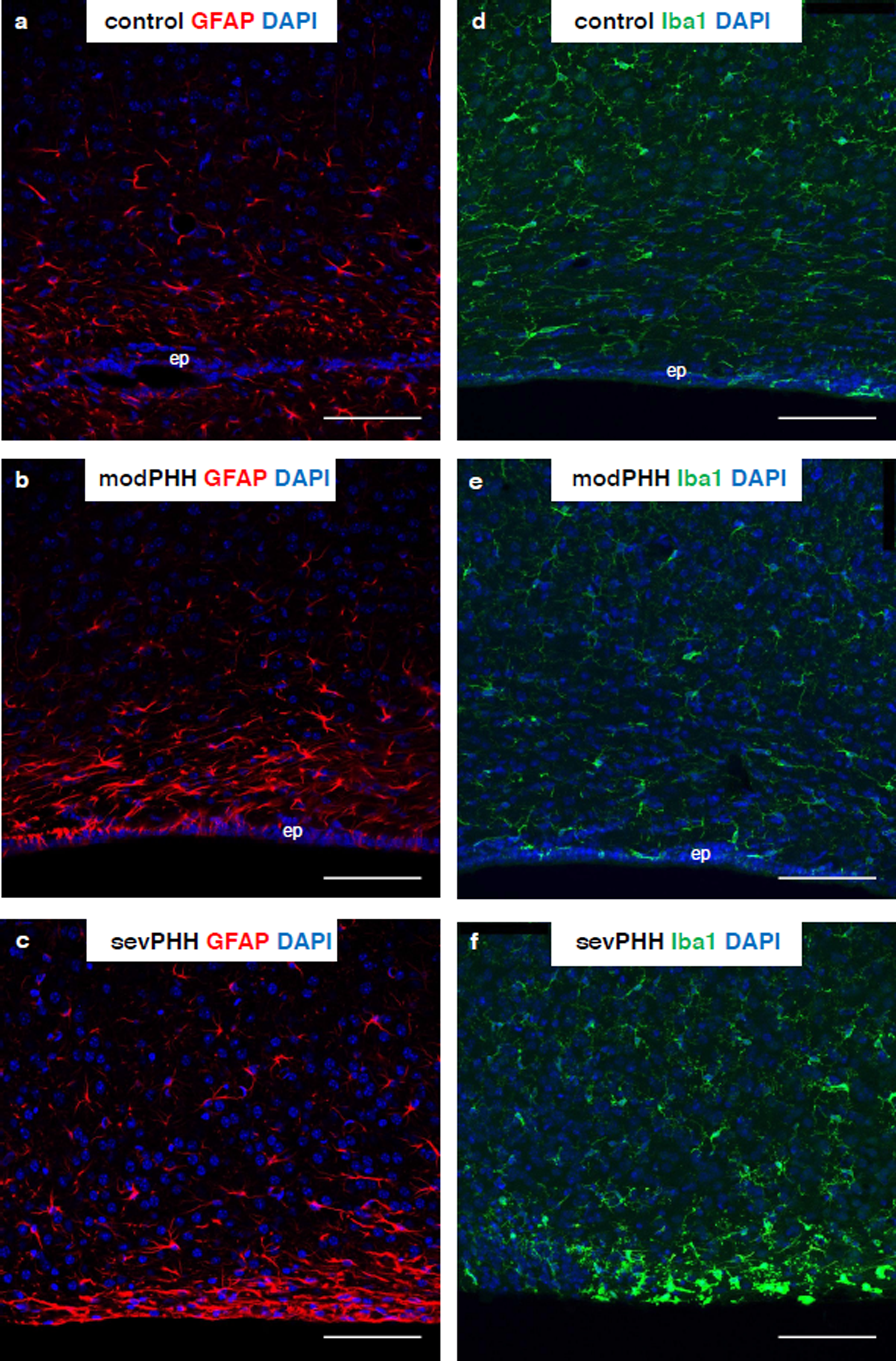
Reactions of astroglia (*GFAP* labeling) and microglia (*Iba1* labeling), compared with a normal (*control*) mouse **(a, d),** in mice with modPHH **(b, e)** and sevPHH **(c, f)**. The presence of ependyma is indicated when present (*ep*). Cell nuclei stained with DAPI (*blue*). Scale bars: 77 µm. The results corresponding to mice with PHH correspond to GMH induction by collagenase injection.

The analysis of overexpressed proteins revealed, in both sevPHH and modPHH, when comparing with controls, alterations in molecular pathways that are indicative of intraventricular hemorrhage consequences, such as the thrombin-activated receptor signaling pathway, fibrinolysis, intracellular sequestration of iron ions, and regulation of blood coagulation (Fig. 3, Tables 1 and 2). In mice with both sevPHH and modPHH, according to overexpressed and undepressed proteins, the presence of astrocyte reactions, defective axonal/myelin development and myelin damage, and cell death were apparent (Tables 1–4). A comparative analysis between the mice with modPHH and sevPHH revealed a differential expression of proteins related to PHH histopathology, including cell processes such as astrocyte reactions, neuronal development, cell death, and myelin damage (Tables 5 and 6). Interestingly, the mice with modPHH showed additional changes in energy metabolism and cell processes related to hypoxia, oxidative stress, and autophagy (Table 4).

**Fig. 3.**
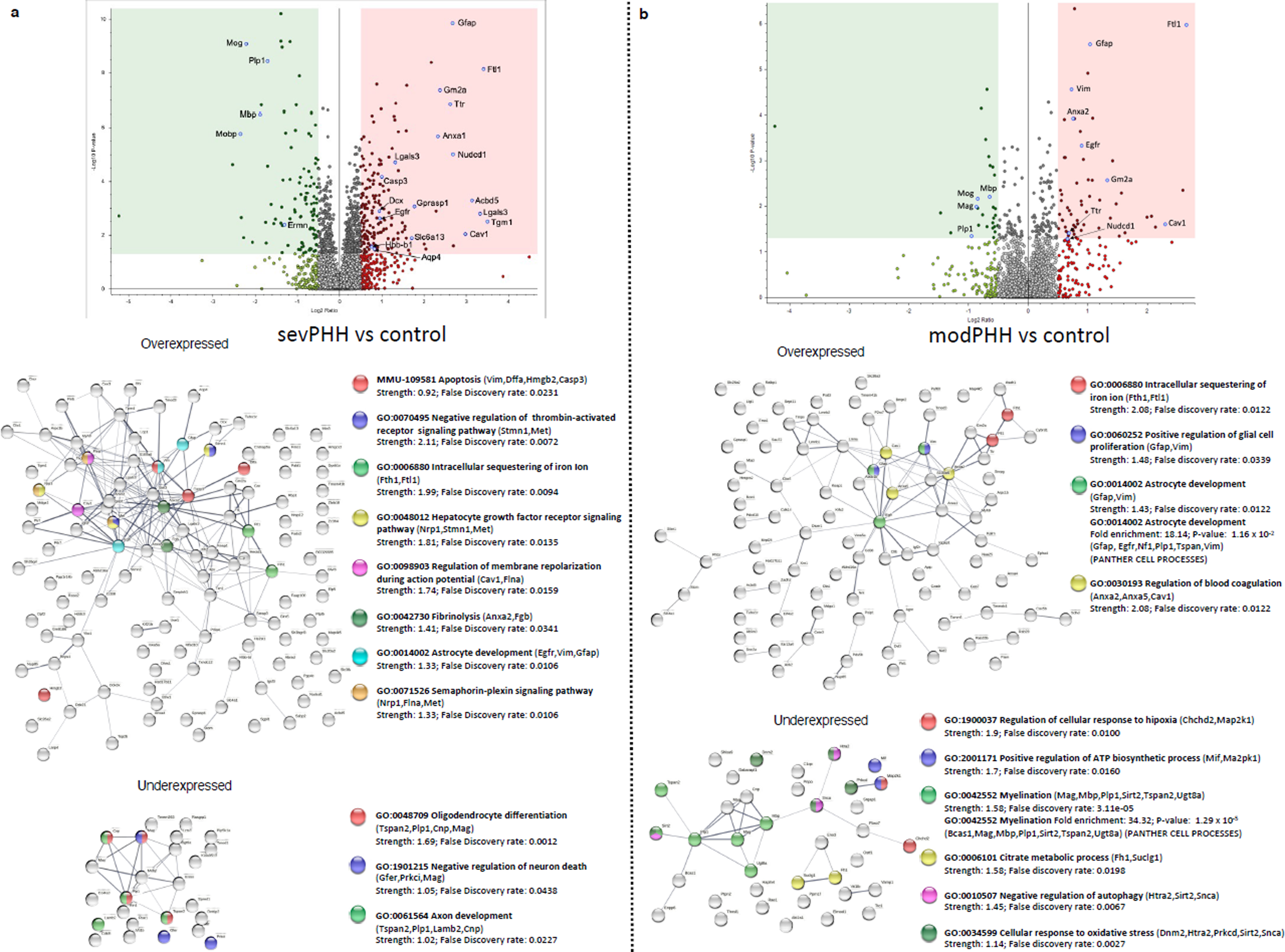
Overexpressed and underexpressed proteins comparing **a** the mice with sevPHH (n = 6) and **b** modPHH (n = 7) with normal mice (*control*, n = 10). Volcano plots (upper), protein‒protein interaction networks and functional enrichment analysis with STRING (bottom) are represented. For details, see the Table 1 legend. Some concordant results from PANTHER GO analyses are also shown and indicated. A fold enrichment indicates that the category was overrepresented. A binomial test with Bonferroni correction for multiple testing with P < 0.05 was used.

**Table 1.** Enhanced cell processes comparing the mice with sevPHH with the controls. The results of GO processes and Reactome pathways (MMU) are shown. Strength represents log10 (observed/expected). This measure describes the size of the enrichment effect. It is the ratio between i) the number of proteins in the network annotated with a term and ii) the number of proteins that are expected to be annotated with this term in a random network of the same size. The false discovery rate describes how significant the enrichment is. The p values corrected for multiple testing within each category using the Benjamini–Hochberg procedure are shown.<colcnt=4>

| GO Process<br>Reactome process (MMU) | Genes | Strength | False<br>Discovery<br>Rate |
| --- | --- | --- | --- |
| GO:0070495 Negative regulation of thrombin-activated receptor signaling pathway | Stmn1, Met | 2.11 | 0.0072 |
| GO:0006880 Intracellular sequestering of iron Ion | Fth1, Ftl1 | 1.99 | 0.0094 |
| GO:0048012 Hepatocyte growth factor receptor signaling pathway | Nrp1, Stmn1, Met | 1.81 | 0.0135 |
| GO:0098903 Regulation of membrane repolarization during action potential | Cav1, Flna | 1.74 | 0.0159 |
| GO:0042730 Fibrinolysis | Anxa2, Fgb | 1.41 | 0.0341 |
| GO:0014002 Astrocyte development | Egfr, Vim, Gfap | 1.33 | 0.0106 |
| GO:0071526 Semaphorin-plexin signaling pathway | Nrp1, Flna, Met | 1.33 | 0.0106 |
| MMU-109581 Apoptosis | Vim, Dffa, Hmgb2, Casp3 | 0.92 | 0.0231 |

**Table 2.** Enhanced cell processes comparing the mice with modPHH with the controls. See legend of Table 1.

| GO Process | Genes | Strength | False<br>Discovery<br>Rate |
| --- | --- | --- | --- |
| GO:0006880 Intracellular<br>sequestering of iron ion | Fth1, Ftl1 | 2.08 | 0.0094 |
| GO:0030193 Regulation of blood<br>coagulation | Anxa2, Anxa5, Cav1 | 2.08 | 0.0122 |
| GO:0060252 Positive regulation of<br>glial cell proliferation | Gfap, Vim | 1.48 | 0.0339 |
| GO:0014002 Astrocyte<br>development | Gfap, Vim | 1.43 | 0.0122 |

**Table 3.** Decreased cell processes comparing the mice with sevPHH with the controls. See legend of Table 1.

| GO Process | Genes | Strength | False<br>Discovery<br>Rate |
| --- | --- | --- | --- |
| GO:0048709<br>Oligodendrocyte<br>differentiation | Tspan2, Plp1, Cnp, Mag | 1.69 | 0.0012 |
| GO:1901215 Negative<br>regulation of neuron death | Gfer, Prkci, Mag | 1.05 | 0.0438 |
| GO:0061564 Axon development | Tspan2, Plp1, Lamb2, Cnp | 1.02 | 0.0227 |

**Table 4.**
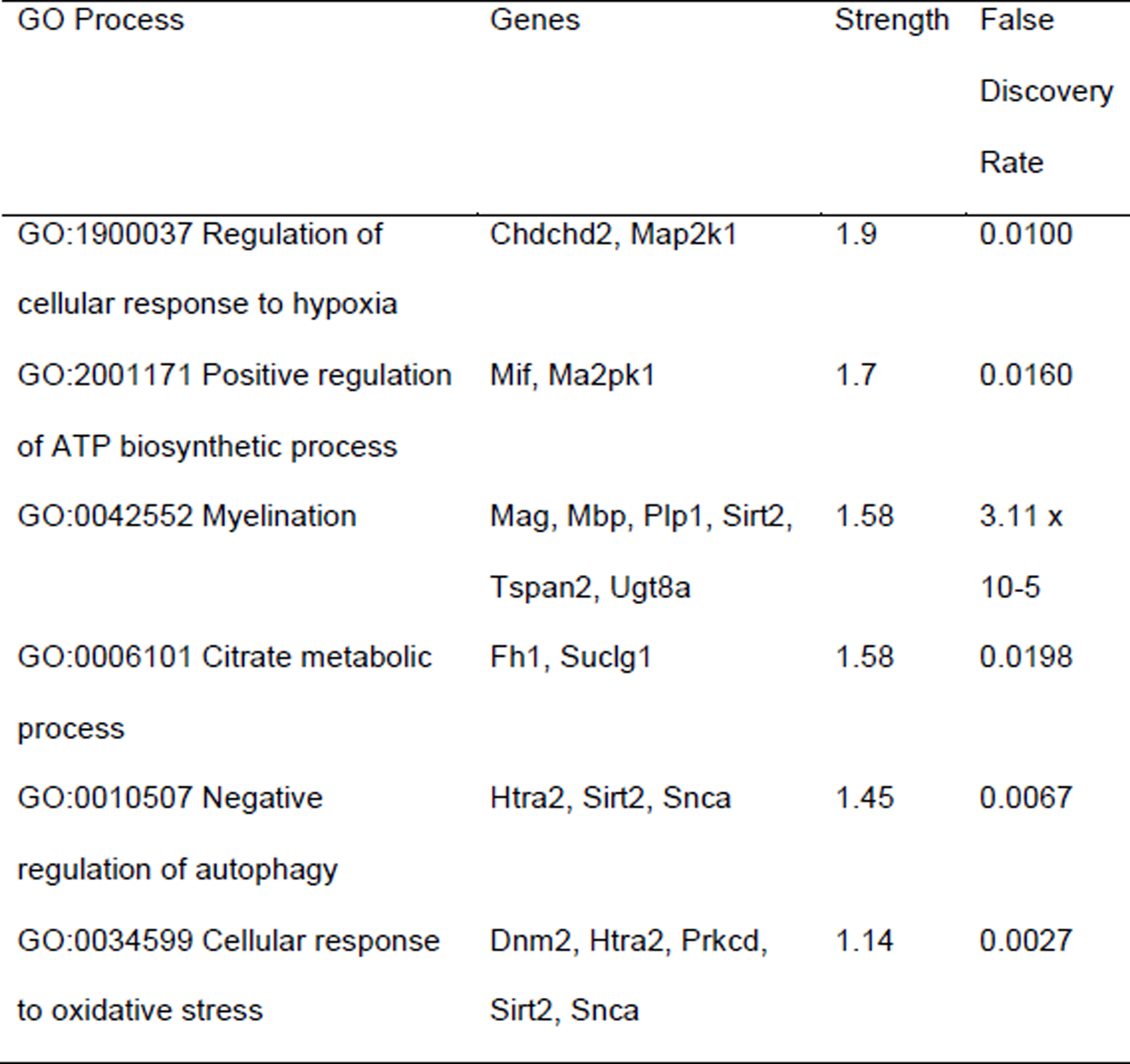
Decreased cell processes comparing the mice with modPHH with the controls. See legend of Table 1.<colcnt=4>

**Table 5.** Enhanced cell processes comparing the mice with sevPHH with those with modPHH. See the legend of Table 1.

| GO Process | Genes | Strength | False<br>Discovery<br>Rate |
| --- | --- | --- | --- |
| Reactome process (MMU) |  |  |  |
| MMU-109581 Apoptosis | Lgals1, Cp | 1.39 | 0.0021 |
| GO:0014002 Astrocyte development | Gfap, Vim | 1.75 | 0.013 |
| GO:0010977 Negative regulation of neuron projection development | Bcl11a, Vim, Flna, Cspg4, Gfap, Lgals1 | 1.46 | 1.78 x 10<sup>-5</sup> |
| GO:0050768 Negative regulation of neurogenesis | Bcl11a, Vim, Flna, Cspg4, Gfap, Lgals1, Dynlt1c | 1.23 | 2.74 x10<sup>-5</sup> |
| GO:0042063 Gliogenesis | Vim, Cspg4, Gfap, Dcx | 1.15 | 0.0045 |

**Table 6.** Decreased cell processes comparing the mice with sevPHH with those with modPHH. See legend of Table 1.

| GO Process | Genes | Strength | False<br>Discovery<br>Rate |
| --- | --- | --- | --- |
| GO:0019911 Structural constituent of myelin sheath | Plp1, Mobp | 2.6 | 0.0012 |
| GO:0017016 Ras GTPase binding | Mobp, Cdkl5, Rasgrp1 | 1.21 | 0.0283 |

### Developing an ex vivo system to test ependymal restoration

Two days after the GMH/IVH was generated with collagenase injection, blood or blood serum, it was impossible to identify the GMH extension nor if the animals were developing moderate (modPHH) or severe (sevPHH) by analyzing external physical signs or by the behavior of the animal. To overcome this problem, a new experimental approach was developed to test regenerative stem cell therapy based on an *ex vivo* system. After analyzing the efficiency of generating GMH/PHH with collagenase, blood or blood serum injections, collagenase and blood serum injections were selected to test ependymal restoration. However, the condition of the whole blood injection was discarded. Collagenase injection showed high efficiency and consistency in generating the GMH, it would mimic blood injection (hematocrit effect as well as intensities of the hemorrhages that were generated), and it would reduce the number of experimental conditions. Blood serum condition was kept as it allowed us to test the effect of the inflammatory factors in the transplanted stem cells and their progeny independently of the blood cellular components. Therefore, three days after collagenase or blood serum intraventricular injection inducing IVH/GMH, animals were sacrificed, ventricular walls exposed and classified into developing moderate (modGMH/PHH) or severe (sevGMH/PHH) according to the GMH extension, ventricular size, and periventricular damage (Fig. 4a, b). Then, walls were dissected out, explants generated and cultured as an *ex vivo* system where the different combinations of stem cell therapies could be tested.

**Fig. 4.**
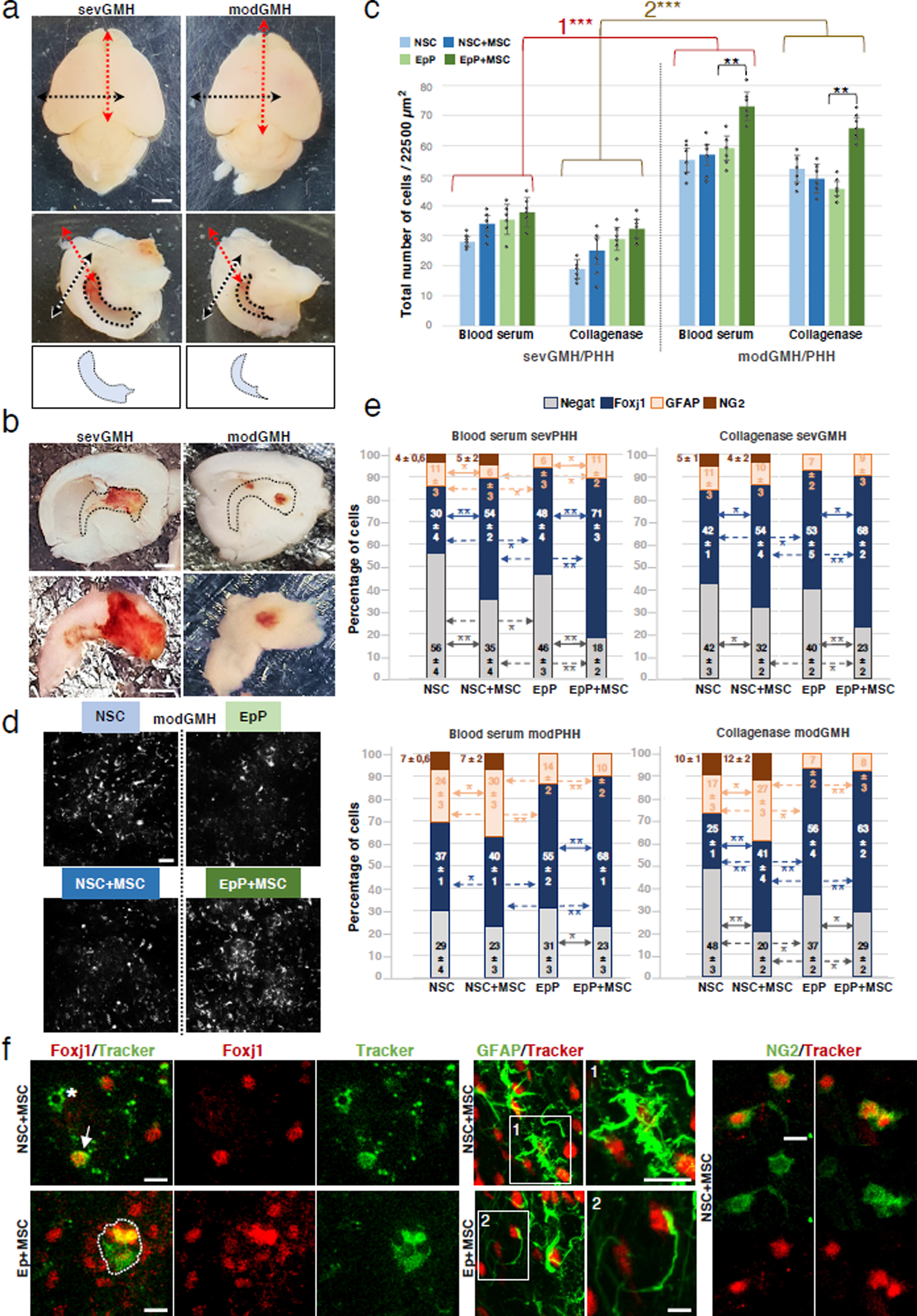
Survival and differentiation of EpPs and NSCs in explants of the ventricular walls from mice that develop modGMH/PHH and sevGMH/PHH after collagenase and blood serum injections. **a** Dissection of ventricular wall explants showing cerebral-ventricular cavities and dilatation with modGMH and sevGMH after collagenase injection. The top pictures show the isolated brain previous dissection. The double-ended red and black dashed lines represent the incision planes that expose cerebral-ventricular cavities. The bottom pictures show differential ventricular dilatation. Perimeters of the cerebral-ventricular cavities are demarked with dotted lines. **b** Exposed lateral ventricular striatal walls showing the periventricular hemorrhages (upper) and dissected explants (bottom). The left pictures correspond to a mouse with sevGMH showing extensive hemorrhage. The right pictures correspond to a mouse with modGMH according to the limited hemorrhage. **c** Survival rates of the NSCs and EpPs transplanted into lateral ventricle explants from the animals with sevGMH/PHH (left) or modGMH/PHH (right) induced by blood serum or collagenase injection. The total number of cells that survived seven days after transplantation is shown. Mean ± SD, n = 5 each condition, ***p < 0.005, **CSp < 0.01, Wilcoxon-Mann-Whitney test. Bars represent standard deviation. Between the mice with modGMH/PHH and sevGMH/PHH with and without MSC-pretreatment, for blood serum (*1*) and collagenase (*2*) conditions, differences in the survival rates of the NSCs and EpPs were significant. **d** Samples of pictures showing the transplanted cells, labeled with a tracker, on the surface of the explant from the animals with modGMH induced by collagenase. **e** Differentiation of the transplanted NSCs and EpPs into Foxj1+ cells, GFAP+ cells, and NG2+ cells seven days after transplantation under different experimental conditions. Transplanted NSCs and EpPs cells negative for these markers are also shown. Mean ± SEM, n = 5 each condition. *p < 0.05, **p < 0.01; Wilcoxon-Mann-Whitney test. **f** Sample pictures used for quantification in ***d*** showing differentiated labeled transplanted cells on the surface of the lateral ventricle explants from animals with modGMH induced by collagenase and under MSC-pretreatment conditions. Left: transplanted NSCs and EpPs (labeled with green cell tracker, *arrow*) expressing Foxj1 in MSC-pretreatment conditions. *Asterisk* points to an NSC lacking expression of Foxj1. A cluster of Foxj1+ cells derived from EpPs is framed. Middle: transplanted NSCs and EpPs (labeled with red cell tracker) expressing GFAP under MSC-pretreatment conditions. Details (*1* and *2*) are shown on the right. Scale bars: **a**, **b**, 2 mm; **d**, 30 µm; **f**, 20 µm.

### Survival rates of transplanted stem cells according to the severity of posthemorrhagic hydrocephalus development

EpP and NSC transplantation was performed in ventricular wall explants of mice that presented modGMH/PHH and sevGMH/PHH to study the possibility of ependymal restoration.

The survival rate of NSCs and EpPs (including their progeny) was studied seven days after transplantation into ventricular wall explants. In addition, the effect of pretreatment with bone marrow-derived MSCs in the explants was analyzed. The survival rate of the transplanted stem cells (NSCs and EpPs) was higher in the explant walls with modGMH/PHH than in those with sevGMH/PHH (Fig. 4c). In addition, pretreatment with MSCs improved the survival rate of EpPs but not NSCs in explants from the mice with modGMH/PHH but had no effect on explants from the mice with sevGMH/PHH (Fig. 4c, d).

### Differentiation of ependymal progenitor cells and neural stem cells into ependymal cells

Once it was proven that the progenitor cells (EpPs and NSCs) could survive into the ventricular wall explants of the mice with modGMH/PHH and sevGMH/PHH (Fig. 4c), their respective efficiencies to produce cells committed to be ependyma expressing the transcription factor Foxj1^50^ was investigated (Fig. 4e).

For description purposes, positive cells derived from EpPs and NSCs are called ^EpP^Foxj1+ and ^NSC^Foxj1+, respectively. A fluorescent cell tracker was added *in vitro* before the application to distinguish the transplanted cells and their progeny. In the lateral ventricle wall explants from the mice showing modGMH/PHH and sevGMH/PHH, the ^EpP^Foxj1+ percentage was higher than the ^NSC^Foxj1+ percentage (Fig. 4e). This result was consistent for blood serum and collagenase injections (Fig. 4e). Therefore, and not unexpectedly, EpPs generated more Foxj1+ cells than NSCs. Notably, MSC-pretreatment in explants from the mice with modGMH/PHH and sevGMH/PHH increased the rates of ^EpP^Foxj1+ and ^NSC^Foxj1+ (Fig. 4e, f).

In addition to Foxj1+ cells derived from NSCs or EpPs, the presence of other main cell types derived from transplanted stem cells was analyzed. For this purpose, the presence of NG2+ and GFAP+ cells and transplanted cells that did not express any of these markers (Foxj1-/NG2-/GFAP-), thus called negative (under immunofluorescence) cells, was also quantified. In the case of transplanted NSCs, other cells expressing NG2 and GFAP were generated (Fig. 4e, f). Transplanted EpPs generated GFAP+ cells but not NG2+ cells under all studied conditions (Fig. 4e, f). Pretreatment with MSCs did not change the type of progeny generated from NSCs or EpPs for the studied markers GFAP and NG2 (Fig. 4e). However, for NSC and EpP transplantation, MSC-pretreatment revealed a generalized decrease in negative cells (Foxj1-/NG2-/GFAP-) that agreed with the increased rate of Foxj1 positive-derived cells (Fig. 4e).

### Cilia development from cells committed to ependymal differentiation under posthemorrhagic hydrocephalus conditions

After it was observed that the transplanted cells (EpPs and NSCs) could generate Foxj1+ cells in the ventricular wall explants of the mice with modGMH/PHH and sevGMH/PHH, the efficiency of these Foxj1+ cells to terminally differentiate into multiciliated ependymal cells was investigated. Cilia were detected with βIV-Tubulin labeling in the Foxj1+ cells in this case. We found that a percentage of Foxj1+ cells did not develop multicilia; therefore, these cells could not fully differentiate into multiciliated ependyma (Fig. 5a, b). In explants from the mice with modGMH/PHH and sevGMH/PHH, higher percentages of ciliated cells were detected when transplanting EpPs compared to NSCs (Fig. 5a, b).

**Fig. 5.**
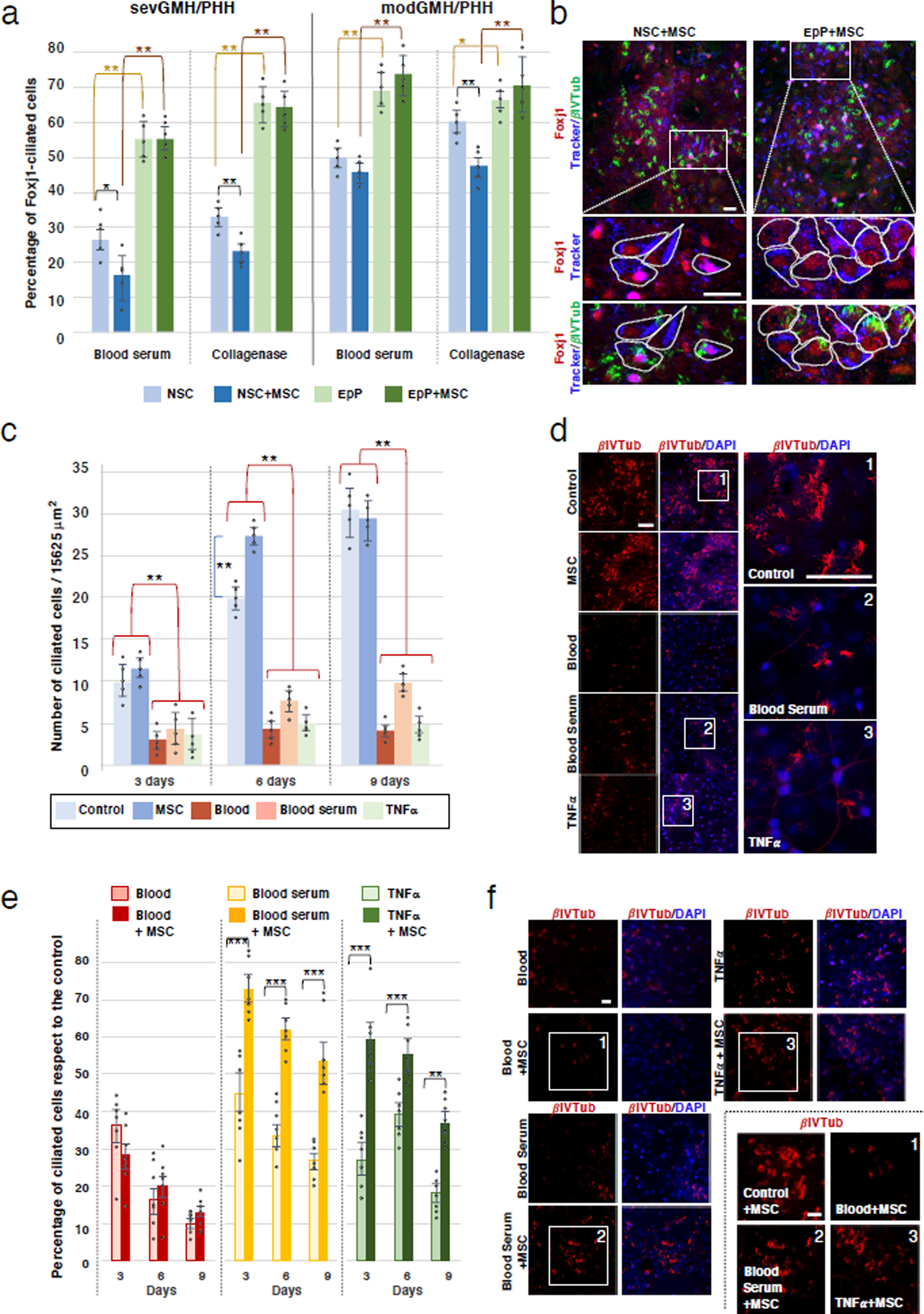
Differentiation into multiciliated ependyma of Foxj1+ cells derived from transplanted NSCs and EpPs. **a** Percentages of ciliated Foxj1+ cells derived from NSCs and EpPs after seven days in ventricular walls of the sevGMH/PHH (left) and mod/GMHPHH groups (right). GMH/PHH was induced by blood serum or collagenase injections. MSC-pretreatment conditions are shown. Mean ± SD, n = 5, *p < 0.05, **p < 0.01; Wilcoxon-Mann-Whitney test. Bars represent the standard deviation. **b** Multiciliated progeny from transplanted EpPs (right) and NSCs (left) after seven days in ventricular wall explants from the animals with modGMH generated by collagenase injection and in MSC-pretreatment conditions. NSC- and EpP-derived cells are labeled with a cell tracker. At the bottom, details framed at the upper are shown. Some of those Foxj1+ cells are multiciliated (cilia labeled with βIV-Tubulin), indicating that they fully differentiate into ependymal cells. **c** Rates from EpPs developing cilia at different times (3, 6, 9 days) in primary cultures and conditions that could be present in GMH/PHH *in vivo:* with blood, blood serum, and TNFα; and in MSC-pretreatment. A control condition is also shown. Mean ± SD, n = 5 each condition. **p < 0.01; ***p < 0.005; Wilcoxon-Mann-Whitney test. Bars represent the standard deviation. **d** Sample pictures used for quantification of cilia development in primary culture of EpPs after nine days in ***c***. Cilia percentage differs under the different culture conditions in ***c***. Detailed images are shown on the corresponding *1*, *2*, and *3* framed areas. **e** Percentages of EpPs developing cilia for the control condition at different times in primary cultures under treatment with blood, blood serum, and TNFα. The effect of MSC-pretreatment is shown for each condition. Mean ± SEM, n = 7 each condition. **p < 0.01, ***p < 0.005; Wilcoxon-Mann-Whitney test. Bars represent the standard error of the mean. **f** Sample pictures of cilia development in primary culture of EpPs after nine days with and without MSC-pretreatment. The percentage of cilia (labeled with βIV-Tubulin) differs under different culture conditions. Detailed images from the *1*, *2*, and *3* framed areas are on the bottom left. Scale bars: **b**, 15 µm; **d**, 30 µm; **f**, 15 µm.

The microenvironmental conditions associated with posthemorrhagic development, including the inflammatory response in the tissue, can affect stem cell differentiation and function^51, 52^. In the explants, pretreatment with MSCs, which can modulate inflammatory conditions, did not increase the percentages of Foxj1+ cells developing cilia from the EpPs (Fig. 5a, b). Surprisingly, the MSC environment decreased the rate of Foxj1+ ciliated cells derived from NSCs (Fig. 5a).

### Effect of treatment with MSCs on cilia development from cells committed to the ependyma under posthemorrhagic hydrocephalus conditions

Due to MSCs did not increase the percentages of Foxj1+ cells developing cilia, we investigated the effect that different elements present in the hemorrhage may have on cilia development *in vitro*. An *in vitro* system was used to avoid the complex inflammatory environment present in the *ex vivo* system. The possible benefit of pretreatment with MSCs was analyzed.

The effect of hemorrhage on ependymal differentiation using primary EpP cell cultures was first studied (Fig. 5c-d). The effect of blood, blood serum, or the inflammatory cytokine TNFα was analyzed. TNFα was selected for its role in hydrocephalus development, astrocytosis, and inflammatory processes^53^. The results revealed that an environment with TNFα, blood or blood serum prevents the development of multicilia in Foxj1+ cells (Fig. 5c, d). This effect was more pronounced in the presence of whole blood and TNFα (Fig. 5c, d). Interestingly, when Foxj1+ cells were cocultured with MSCs under control conditions (noninflammatory environment), the development of multicilia was hastened (Fig. 5c). However, this MSC modulation did not affect the final percentage of differentiated multiciliated ependymal cells (Fig. 5c, d).

Finally, the possible therapeutic effect of MSCs on the ciliogenesis of cells derived from EpPs after exposure to TNFα, blood or blood serum was studied. Coculture of EpPs and MSCs was performed following EpP exposure to TNFα, blood or blood serum, and Foxj1+ cells presenting cilia were quantified. Interestingly, MSC coculture improved the percentages of Foxj1+ cells developing cilia in the presence of blood serum or TNFα but not whole blood (Fig. 5e, f).

### Effects of stem cell therapy on the brain parenchyma

Once that MSC effect on ciliogenesis was analyzed *in vitro* and its positive role in helping ciliogenesis under inflammatory conditions was observed, the impact on brain parenchyma was studied using our *ex vivo* ventricular wall explants system after collagenase injection. For this experiment, the optimal condition, e.g., EpP transplantation into collagenase-induced modGMH, was selected based on the results of previous experiments (Figs. 4c, 4e, and 5a).

Ventricular walls were dissected 48 hours after GMH induction, and MSCs were added to the explant. Twenty-four hours later, EpPs were also transplanted, and the explants were kept in these culture conditions for seven days. Evans Blue was applied after treatment, and the tissue was fixed and analyzed under confocal microscopy. The staining intensity indicated cell damage and edematous grade status; more intensity was equivalent to major edema. The intensity of the staining/edema was classified into three different categories to evaluate the possible effect of the treatment. These categories were created based in the intensity of the staining: severe edema, corresponding to 70-100% of staining intensity; moderate edema, 25-69% of staining intensity; and light edema, 0-24% of staining intensity. The percentage of surface presenting severe, moderate, or light edema was analyzed and compared per experimental condition (Fig. 6a). In control conditions, without MSCs, severe edema represented 18% of the ventricular surface, and moderate edema represented 67% (Fig. 6a, b). Transplantation of MSCs revealed a surface reduction of severe edema to 2% and moderate edema to 14% (Fig. 6a). However, the area less affected by edema increased from 14% to 84% (Fig. 6a, b). When both MSCs and EpPs were transplanted into ventricular explants, the surfaces with severe and moderate edema decreased (Fig. 6a, b). Conversely, the area less affected by edema with treatment increased (Fig. 6a, b).

**Fig. 6.**
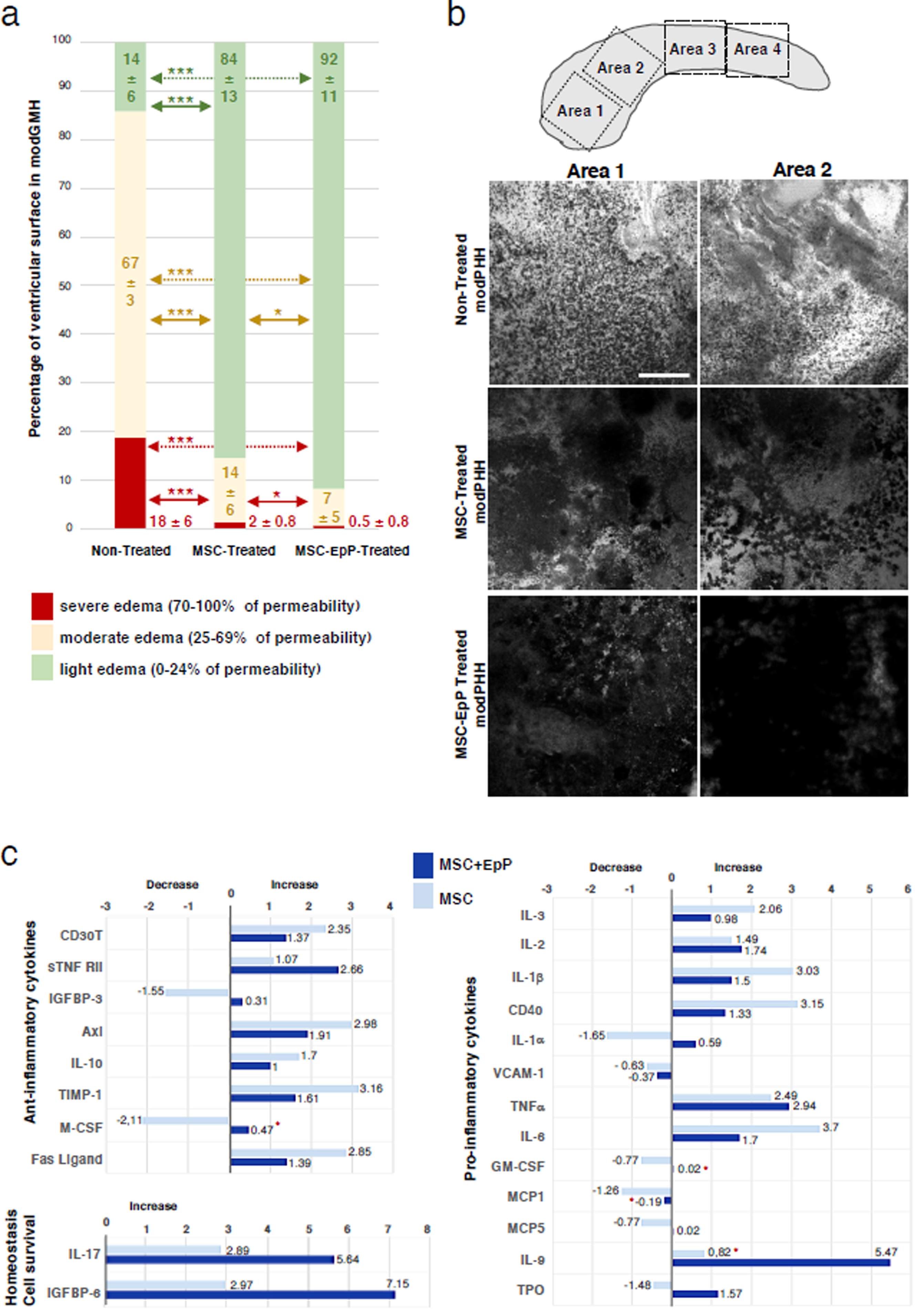
Effect on brain parenchyma after transplantation with EpPs and MSCs. **a** Percentages of ventricle extensions displaying different degrees of damage stained with Evans Blue. The 70-100% highest intensity for Evans blue staining was considered severe edema, 25-69% moderate edema, and 0-24% light edema. Mean ± SD, n = 6, *p < 0.05, ***p < 0.005; Wilcoxon-Mann-Whitney test. **b** Representative ventricle surface images from explants stained with Evans Blue from the mice treated with modHMH and sevGMH seven days after treatment with EpPs and MSCs. The images correspond to framed areas *1* and *2* of the explant *in vitro* drawn on the upper part. **c** Fold changes compared with the control condition in cytokine content obtained by comparing MSC-pretreated explant medium versus nontreated explant medium and MSC+EpP-treated explant medium versus nontreated explant medium. The cytokines are grouped as anti-inflammatory cytokines, favoring cell survival and homeostasis, and pro-inflammatory cytokines. Positive values indicate an increase versus nontreated explants, while negative values indicate a decrease versus nontreated explants. Asterisks show nonsignificant differences in the values of cytokine content in medium between stem cell treated explants versus nontreated explants. Scale bar: **b**, 250 µm.

### Effects of stem cell therapy on the cytokines present in the explant media

From the experiments performed to analyze the effects of stem cell therapy on the brain parenchyma described above, culture media were collected to detect cytokines under the different treatments: nontreated explants, MSC-pretreated explants, and MSC+EpP-treated explants. Cell culture medium without explants was used as a baseline control. The analysis of the 62 cytokines analyzed focused on those that presented significant differences with the nontreated explants (Fig. 6c). Fold changes were registered in the expression of the cytokines between the MSC-pretreated explants and MSC+EpP-treated explants versus the nontreated explants (Fig. 6c).

The levels of anti-inflammatory cytokines were generally increased in the MSC-pretreated explants compared to the nontreated explants, except for IGFBP-3 and M-CSF (Fig. 6c). Cytokines related to cell survival and homeostasis, such as IL17 and IGFBP-6, increased in the MSC-pretreated versus non-pretreated explants (Fig. 6C).

In the analysis of pro-inflammatory cytokines, the results were more variable (Fig. 6c). The media were collected at the end of the experiment; therefore, a general increase in pro-inflammatory cytokines generated from the first hours after transplantation could be expected. For some pro-inflammatory cytokines, such an increase was lower in the explants with EpP-treatment (Fig. 6c).

## Discussion

Ependyma is a relevant structure for brain development, homeostasis, and neurogenesis. This structure is severely damaged after GMH/PHH. However, treatments for GMH/PHH are not directed to treat the ependyma. Consequently, severe neurological defects associated with ependyma alteration persist lifelong with almost no improvement in patient outcomes.

The present investigation is directed to generate new therapeutic options, clinically relevant to overcome the ependymal gap when treating GMH/PHH. In this way, the present study has tested the possibility of repairing the ependyma in PHH to help tissue recovery and prevent pathological conditions associated with PHH and ependyma damage, such as periventricular edema. For this purpose, a PHH animal model that mimics the neuropathological events occurring in human cases, including ependymal affectation, has been developed. Additionally, a new *ex vivo* system based on ventricular wall explants affected with induced GMH/PHH has been designed to test the possibility of ependymal restoration. Finally, NSCs or EpPs were transplanted, their ependymal progeny was identified and quantified, the effect of pretreatment with MSCs was analyzed, and their positive impact on tissue recovery was proven.

### Pathological processes related to GMH/PHH severity indicate the relevance of ependymal restoration

Several animal models have been created to develop PHH experimentally^54, 55^. A series of aspects related to the state of maturity of the periventricular tissues have been proposed as relevant for experimentally studying the development of PHH^56^. The present investigation produced PHHs according to the three most common experimental procedures^57^. In all three, common cytopathology has been found consisting of ependymal disruption associated with reactions of microglia and astroglia cells. Notably, the degree of those alterations correlates with the severity of hydrocephalus, which the molecular analysis has corroborated. The results on differential protein expression indicate that the PHH pathogenesis present in the modPHH and sevPHH cases can be grouped. The first group comprises several blood components, such as hemoglobin, iron, and thrombin, which are implicated in PHH pathogenesis and defects in brain development^35, 57–60^. A second group indicates astrocytic activation following IVH and subsequent inflammation, which is a primary determinant of white matter brain damage in preterm infants^6, 49, 59^. Microglial and astrocytic activation following GMH/IVH and subsequent inflammation are primary determinants of white matter brain damage in preterm infants^61^. A third group could be constituted by proteins implicated in cell death and axonal injury in the periventricular cerebral parenchyma^15, 62, 63^. The presence of blood components has been proven to be directly related to the disruption of ependymal development by affecting cell junctions and activating the periventricular astroglial reaction^16, 64^. Lysophosphatidic acid in the blood appears to be implicated in ependymal disruption^26, 27^. Ependymal disruption can be associated with the development of a periventricular astrocyte reaction, which has also been shown in human cases^12^. In PHH, this reaction is present at the ventricle surface and in the white matter, similar to other forms of fetal-neonatal hydrocephalus^21, 65, 66^.

### A combination of mesenchymal stem cells and ependymal progenitors appears to be an adequate therapeutic strategy

The results of the present investigation indicate that repair of the ependyma from NSCs and EpPs is possible in GMH/PHH pathology.

Inflammation and microglial activation have been proven to be key events in the different forms of congenital and acquired fetal-neonatal hydrocephalus, including those of GMH/IVH origin^13, 31, 66–70^. In the present investigation, the observed effects of MSC-pretreatment can be related to their immunomodulatory properties^71, 72^.

In premature children with GMH/PHH, NSCs have been obtained from CSF collected during neuroendoscopic lavage^73^. These NSCs could therefore be used for regenerative purposes^73^. Interestingly, neuroendoscopic lavage can remove blood-derived components, attenuate inflammatory conditions, and thus favor ependymal differentiation. However, as shown in the present results, our experimental approach shows that using EpPs would be more successful in restoring ependyma.

Identifying the most efficient stem cell combination for ependymal production and survival under these inflammatory conditions was crucial to study a possible stem cell therapeutic strategy for ependymal restoration after GMH/PHH. The results of the present investigation have revealed that MSCs influence transplanted EpP survival. Mesenchymal stem cells have immunomodulatory and neuroprotective effects^74, 75^, which make them suitable for treating CNS diseases mediated by the immune system, such as neurodegenerative diseases^76^ or hydrocephalus^43^. The neuroprotective role of MSCs is based on creating favorable environments for regeneration, producing growth factors and cytokines, and promoting vascularization and myelination^74^. The secretion of anti-inflammatory molecules and modulation of specific molecular pathways in the EpP differentiation process could stabilize them under adverse conditions.

The results of the present investigation also indicate that EpPs have a better ability to generate ependyma than NSCs under inflammatory conditions present in GMH/PHH. This difference between NSCs and EpPs became more evident when these two populations were transplanted after pretreatment with MSCs. In normal conditions, the starting potential of NSCs and EpPs is different and could be key for understanding the present results and their therapeutic implications. Although NSCs display the potential for neuron, oligodendrocyte and ependymal differentiation *in vitro*, it has been reported that most NSCs differentiate into astrocytes when transplanted *in vivo* under inflammatory conditions^77^. Under inflammatory conditions, proinflammatory signals such as TNFα or BMPs can alter the expected behavior of NSCs, modifying their proliferation rate and normal differentiation process and reducing the type and amount of their progeny^78, 79^. However, although the transplanted EpPs will be most committed to differentiating into ependymal cells, proinflammatory cytokines can induce the loss of mature ependymal status^80, 81^. Both NSCs and EpPs produced more Foxj1+ cells after MSC-pretreatment, thus indicating that MSCs generate a more favorable environment for NSCs and EpPs to give rise to the cell types preprogrammed to produce. Thus, the EpPs that survive after transplantation have been shown to give rise primarily to one cell type, Foxj1+ cells. In contrast, NSCs were found to produce different cell types; only a proportion of their progeny were Foxj1+ cells. Therefore, the EpPs were the more appropriate cells for ependymal restoration after GMH/PHH, and MSC-pretreatment would be favorable. Under pro- and anti-inflammatory conditions, stem cells can modify their progeny^78, 79^. However, our results showed that after MSC-pretreatment, there was no change in the type of progeny generated from NSCs and EpPs for the studied markers.

### TNFα and blood components affect the development of ependymal cilia

In the present work, cells derived from EpPs expressing Foxj1 showed alterations in ciliogenesis in experiments with ventricular wall explants and primary cultures in the presence of an inflammatory environment, blood components, and TNFα.

An inflammatory response has been demonstrated to be present after experimental and clinical GMH/IVH^82–84^. Elevations in inflammatory markers in blood and serum following spontaneous IVH have been reported in clinical studies^85–89^. Blood degradation products activate cytokine production and immune system cell recruitment^90–94^. Blood products related to inflammatory signaling pathways can activate immune cell types and, interestingly, other cell types, including the ependyma^95–98^. TNFα is one of the key molecules expressed in blood or blood serum after IVH^89^, and it has also been found to be expressed in ependyma under injury, ischemia or infection^99, 100^.

Cilia are key ependymal structures present in the ependyma mature stage and implied in CSF circulation^17^. In the present study, the presence of TNFα and blood affected cilia development from Foxj1+ cells. Normal ependymal differentiation requires the Foxj1 transcription factor^101^, and a differentiated multiciliated ependymal state requires constant maintenance of Foxj1^30^. The Foxj1 transcription factor has a short half-life; the ubiquitin-proteasome system rapidly turns it over. Consequently, when Foxj1 degradation occurs, multicilia are not maintained, and ependymal cells dedifferentiate^30^. In Foxj1 transcription factor maintenance, inhibitor kappa β kinase 2 (IKK2) activity is crucial. Thus, IKK2 inhibitors, including viruses and growth factors, robustly induce Foxj1 degradation, ependymal cell dedifferentiation, and hydrocephalus^30^.

In addition, pretreatment with MSCs on the ependyma could reduce the negative effect of TNFα on ciliogenesis. Previous reports have shown that MSCs, under the exposure of high TNFα levels, activate the anti-inflammatory pathway mediated by its receptor TNFR2^102^ and induce their secretion of anti-inflammatory molecules such as TGFβ and IL-10^103^. Interestingly, IL10 is one of the anti-inflammatory cytokines that significantly increases in explants from modGMH mice treated with MSCs. The possible anti-inflammatory pathways activated by MSCs could be numerous, and many of them could converge in the recovery of the ependyma. However, the result would be the stabilization of Foxj1 and, therefore, should recover IKK as a stabilizer. An additional pathway that could be involved in recovery is through Toll-like receptors (TLRs). Blood degradation products bind to TLR4 and activate the IKK-NF-κB pathway as a pro-inflammatory pathway^95–98^. Ependymal cells physiologically express TLR4^99, 104^, which induces activation of the IKK-NF-κB pathway^105^. IL10 secreted by MSCs under inflammatory conditions (including TNFα) can cause TLR4 inhibition^106^. Therefore, MSCs could be helping to recover ependymal cilia and Foxj1 transcription factor maintenance by secreting anti-inflammatory cytokines such as IL10 that could help stabilize the Foxj1 transcription factor activating anti-inflammatory pathways in the ependyma, similar to those controlled by TLR4. Therefore, several molecular mechanisms can explain the present results where MSC activation by pro-inflammatory molecules would block the inflammatory pathways that prevent ependyma from stabilizing its ciliated state. Some of them could be deeply studied for therapeutic purposes.

### Treatment with mesenchymal stem cells and ependymal progenitor cells improves edema conditions in explants

The results of the present investigation indicate that transplantation of MSCs can reduce the severity of edema located on the surface of lateral wall ventricle explants. Therefore, MSC transplantation was either helping the recovery of the damaged ventricular wall or preventing ventricular wall degeneration after GMH/PHH.

As described above, GMH/IVH induces an elevation of proinflammatory cytokines associated with edema in the brain parenchyma^9, 10^. MSCs have a main anti-inflammatory role and an effect on edema reduction under different pathological conditions^43, 74–76^. Accordingly, our results showed that MSC-pretreatment generated a better environment for cell therapy.

Although the positive effect of MSC-pretreatment on the ventricular wall surface was evident, interestingly, this effect was potentiated after EpP transplantation and consequent ependymal restoration. These results may have a promising therapeutic value. As we have discussed, we know that pretreatment with MSCs potentiates ependymal survival generated from EpP transplantation. It should also be considered that ependyma constitutes an essential barrier between CSF and parenchyma and modulates stem cell populations^17–19, 65^. Our results regarding the sequential transplantation of MSCs before neural stem cells suggest a clinical strategy to be considered for any cell therapy in GMH/PHH. Moreover, the use of MSCs can be substituted by their secretion or extracellular vesicles^107, 108^.

## Conclusions

In GMH/PHH, ependyma can be restored by NSCs and more efficiently reestablished by EpPs. This restoration is more effective when sequential stem cell therapy is used with a MSCs administered before the transplantation of NSCs and EpPs. The inflammatory conditions appear to be determinants for survival and differentiation of the transplanted progenitors, where MSC-pretreatment is favorable. The presence of blood components and proinflammatory molecules such as TNFα is responsible for failures in the final differentiation of Foxj1+ cells toward multiciliated ependymal cells, and this effect could be recovered, at least partially, by cotreatment with MSCs.

With actual treatments, severe neurological damage associated with GMH/PHH persists with a slight improvement in patient outcome. Therefore, the presented sequential stem cell therapy using MSCs and EpPs seems to be a promising strategy, clinically relevant to achieve ependymal repair and tissue recovery from neuropathological and neuroinflammatory conditions associated with GMH/PHH.

## Acknowledgments

The authors wish to thank David Navas, Jessica Román, and Remedios Crespillo from the Microscopy, Proteomic, and Molecular Biology Services of the University of Malaga (Spain) for their valuable technical support and all the staff of the Animal Experimentation Service of the University of Malaga (Spain) for their support during the experiments. MRI studies were performed at the ICTS “NANBIOSIS”, specifically in Unit 28 at the “Instituto de Investigación Biomédica de Málaga y Plataforma en Nanomedicina (IBIMA Plataforma BIONAND)”. The present work was supported by grant PI19/00778 (to AJJ and PP-G) from the Instituto de Salud Carlos III, Spain, cofinanced by FEDER funds from the European Union; FPU13/02906 to MG-B from the Ministerio de Educación, Cultura y Deporte, Spain. RYC-2014-16980 to PP-G from the Ministerio de Economía y Competitividad, Spain; UMA18-FEDERJA-277 from Plan Operativo FEDER Andalucía 2014-2020 and Universidad de Málaga to PP-G; Contrato Postdoctoral-PPITD-UMA from Universidad de Málaga to LMR-P; and Proyectos dirigidos por jóvenes investigadores from Universidad de Málaga to PP-G.

## Contribution of authors

Conceptualization and experimental design, AJJ, PPG; generation and characterization of the experimental model of hydrocephaluy, AJJ, BOP, MGM, DDP, CCG; MSC culture, BO, MGB, MGG; obtention of stem cells, BFM, LMRP, PPG; generation of explants, PPG; trasplant of stem cells, PPG; quantification of data, JLS, LMRP; analysys of results, AJJ, PPG; discussion and correction of the manuscript, all the authors; writing the manuscript, AJJ, PPG.

## Availability of data and materials

The datasets used and/or analyzed during the current study are present in the paper and are available from the corresponding authors.

## Ethics approval

The design of the experiments, housing, handling, care, and processing of the animals were conducted following European and Spanish laws (RD53/2013 and 2010/63UE) and ARRIVE guidelines. According to current legislation, experimental procedures (protocol 23/04/2019/069) were approved by the Institutional Animal Care and Use Committee of the University of Malaga, Spain (CEUMA) and the Regional Government Council (Junta de Andalucía, Spain).

## Methods

### Experimental animals

The design of the experiments and animal housing, handling, care, and processing were conducted following European and Spanish laws (RD53/2013 and 2010/63UE) and ARRIVE guidelines^109^. According to current legislation, experimental procedures (protocol 23/04/2019/069) were approved by the Institutional Animal Care and Use Committee of the University of Malaga (CEUMA, Spain) and the Regional Government Council (Junta de Andalucía, Spain).

Mice (C57/BL-6J strain) were originally obtained from Charles River Laboratory and bred in the Animal Experimentation Service of the University of Malaga at 22 °C with a 12:12 light/dark cycle and standard food and water available *ad libitum*.

### GVH/PHH induction

Four-day-old male and female mice were anesthetized with 2% isoflurane in 0.5 l/minute oxygen anesthesia and injected with blood serum (9 µl in the right lateral ventricle or both right and left lateral ventricles), whole blood (4 µl in both right and left lateral ventricles), or 1 µl of sterile saline 0.9% NaCl containing 0.05 U/µl collagenase I (C2674-1G, Sigma‒ Aldrich, St Louis, MO, USA) in the subventricular germinal matrix of each hemisphere, to produce GMH/IVH conditions. The injected blood serum and whole blood were obtained from littermates after decapitation. Blood serum was obtained as the supernatant after centrifugation at 1,400 g for 10 minutes at 4 °C. Blood serum and whole blood injections were performed free-hand using a 26-gauge syringe (Hamilton Syringe 75N, Hamilton, Reno, NV, USA) at coordinates 1 mm posterior to the eye, 1 mm superior to the orbit, and 1 mm deep. Collagenase was injected free-hand using a 33-gauge syringe (Hamilton Neuros, HAMI65460-03) at coordinates 0.5 mm posterior to the eye, 1 mm superior to the orbit, and 2 mm deep. The coordinates were previously assayed with trypan blue injections. The aim of the collagenase injections in the subventricular GM was to mimic premature human GMH/IVH. Fourteen days after the injections, mice were sacrificed, and brains were processed to detect the main cytopathologies associated with PHH development. Normal littermates of the same age without any surgical procedure were used as controls.

### Immunofluorescence in brain sections

Eighteen-day-old male and female mice from the different experimental groups (blood serum in the right lateral ventricle; n = 7; blood serum in both lateral ventricles, n = 4; whole blood, n = 7; collagenase, n = 9; controls without any injection, n = 9; controls with saline serum injection, n = 6) were anesthetized with Dolethal (sodium pentobarbital; Vétoquinol, Lure, France; intraperitoneal administration, 0.2 mg/g body weight) and transcardially perfused with 4% paraformaldehyde diluted in 0.1 M, pH 7.2, phosphate buffer (PB). Fixed brains were dissected out and postfixed in the same solution for 24 hours at 4 °C. Then, brains were cryoprotected in 30% sucrose to obtain frozen sections (60 μm thick). Sections were processed with a free-floating section staining protocol for immunofluorescence using unconjugated primary antibodies (βIV-Tubulin, T7941, Sigma‒Aldrich, dilution 1:400; GFAP, glial fibrillary acidic protein, G-A-5, Sigma‒Aldrich, 1:1000; and Iba1, 019-19741, Wako, Chuoku, Osaka, Japan, 1:500). Secondary antibodies conjugated with Alexa Fluor 488 or Alexa Fluor 568 (Thermo Fisher, Waltham, MA, USA) were used. The antibodies were diluted in PB saline (PBS) containing 0.05% Triton X-100, 0.01% sodium azide, 1% bovine albumin (05477, Sigma‒Aldrich), and 5% appropriate normal sera. Primary antibody incubations were performed for 18 hours at 22 °C or 72 hours at 4 °C. Secondary antibody incubations were performed for 1 hour at 22 °C. Nuclei staining was performed with 4’,6 diamidino-2-phenylindole dihydrochloride (DAPI, Molecular Probes, Eugene, OR, USA) at 300 nM in PBS. For negative controls, the primary antibodies were omitted.

### Magnetic resonance imaging

Magnetic resonance imaging (MRI) experiments were performed on a 9.4T Bruker Biospec small animal MRI system (Bruker Biospec, Bruker BioSpin, Ettlingen, Germany) equipped with a 40 mm quadrature bird-cage resonator and 440 mT/m gradients. Normal male and female mice (n = 5) and mice exhibiting modPHH (n = 3) and sevPHH (n = 2) were anesthetized with 1% isoflurane in 1 l/min oxygen, and body temperature and respiratory frequency were monitored throughout the experiment. The acquisition protocol consisted of a high-resolution T2-weighted RARE (Rapid Acquisition with Relaxation Enhancement) sequence with fat suppression and 3D FISP (Fast Imaging with Steady Precession). Sequence parameters were set as follows. T2-weighted: TR = 2.5 s; effective TE = 32 ms; echo train length = 6; 4 averages; matrix size = 512 × 512; FOV = 20 × 20 mm2; slice thickness = 0.75 mm. 3D FISP: TR = 8 ms, TE = 4 ms, matrix size = 256 × 256 × 128; resolution = 78 μm × 78 μm × 156 μm.

Lateral ventricle size was calculated from high-resolution T2-weighted images as the sum of the volumes (area x slice thickness) of contiguous slices using FIJI 1.53q software (NIH, USA)^110^.

### UHPLC-HRMS analysis of protein expression

For ultrahigh-performance liquid chromatography high-resolution mass spectrometry (UHPLC-HRMS), the caudal cerebral wall was dissected from control mice (n = 10) and mice with modPHH (n = 7) and sevPHH (n = 6) at 18 days of age. Immediately after sacrifice by cervical dislocation, the tissue was frozen on dry ice and stored at −80 °C until further processing. The complete procedure took 2-5 minutes.

Proteins from the samples were purified by a modified trichloroacetic acid protein precipitation procedure (Clean-Up Kit; GE Healthcare, Munich, Germany), and gel-assisted proteolysis was carried out. Briefly, the protein solution was entrapped in a polyacrylamide gel matrix before reduction with dithiothreitol and cysteine residue carbamidomethylation with iodoacetamide. Then, the proteins were digested by trypsin (Promega, Madison, WI, USA), and peptides were extracted from the gel with an acetonitrile/formic acid solution. After extraction, the peptides were purified and concentrated using a C18 ZipTip (Merck Millipore) according to the manufacturer’s instructions.

Samples were injected into an Easy nLC 1200 UHPLC system coupled to a Q Exactive™ HF-X Hybrid Quadrupole-Orbitrap mass spectrometer (Thermo Fisher). Data were acquired using Tune 2.9 and Xcalibur 4.1.31.9 (Thermo Fisher). Peptides from the samples were automatically loaded into a trap column (Acclaim PepMap 100, 75 μm Å∼ 2 cm, C18, 3 μm, 100 A, Thermo Fisher) and eluted onto a 50 cm analytical column (PepMap RSLC C18, 2 μm, 100 A, 75 μm Å∼ 50 cm, Thermo Fisher). The binary gradient mobile phase consisted of 0.1% formic acid in water (solvent A) and 0.1% formic acid in 80% acetonitrile (solvent B). Peptides were eluted from the analytical column with a 120-minute gradient from 2 to 20% solvent B, followed by a 30-minute gradient from 20 to 35% solvent B and finally 95% solvent B for 15 minutes before re-equilibration in 2% solvent B at a constant flow rate of 300 nL/minute. Data acquisition was performed in electrospray ionization positive mode. MS1 scans were acquired from m/z 300–1750 at a resolution of 120000. With a data-dependent acquisition method, the 20 most intense precursor ions with + 2 to + 5 charges were isolated within a 1.2 m/z window and fragmented to obtain the corresponding MS/MS spectra. The fragment ions were generated in a higher-energy collisional dissociation cell with a fixed first mass at 110 m/z and detected by the Orbitrap mass analyzer at a resolution of 30000.

The raw data were analyzed using Proteome Discoverer^TM^ 2.4 (Thermo Fisher). For the identification of the MS2 spectra, Sequest HT was utilized as the search engine, and the Swiss-Prot part of UniProt for *Mus musculus* was utilized as the database. Protein assignments were validated using the Percolator algorithm^111^ by imposing a strict 1% false discovery rate (FDR) cutoff.

Label-free quantitation was implemented using the Minora feature of Proteome Discoverer^TM^ 2.4. Protein abundance ratios were directly calculated from the grouped protein abundances. ANOVA was based on the abundance of individual proteins or peptides.

The PANTHER Classification System (v.14.1)^112^ and GO Ontology database^113^ were used to identify the main biological processes related to the overexpressed/underexpressed proteins. Additionally, the STRING platform was used for functional enrichment analysis with the representation of protein‒protein interaction networks^114^.

### Ventricle wall explant in vitro assays

Forty-eight hours after GMH/PHH induction with collagenase or blood serum, mice were sacrificed by decapitation, and the brain was dissected out and classified according to GMH extension and lateral ventricle size (in collagenase, blood serum and whole blood injections) in modGMH/PHH and sevGMH/PHH. Explants from the lateral wall of the lateral ventricle, the striatal wall, were carefully positioned on Millicell culture inserts (PICM0RG50, Sigma‒Aldrich) into six-well culture plates (three explants/insert) containing 1 mL of sterile organotypic culture medium (26.6 mM HEPES, pH 7.1; 19.3 mM NaCl; 5 mM NaHCO_3_; 511 μM ascorbic acid; 40 mM glucose; 2.7 mM CaCl_2_; 2.5 mM MgSO_4_; 0.033% v/v insulin, I6634, Sigma‒Aldrich; and 0.5% v/v penicillin/streptomycin, P0781, Sigma‒ Aldrich) in ultrapure H_2_O, sterile filtered, with 25% (v/v) heat-inactivated horse serum (H1138, Sigma‒Aldrich). Explants were incubated at 37 °C, and the culture medium was partially replaced every 48 hours. Explants were maintained for a maximum of 7 days *in vitro* before harvesting.

The explants used to study the survival of transplanted stem cells and their progenies were fixed with 4% paraformaldehyde in PB 0.1 M, pH 7.2 at 4 °C for 24 hours and then immunostained using unconjugated primary antibodies followed by secondary antibodies conjugated with fluorochromes, as described above for brain sections. In addition to anti-GFAP and anti-βIVTubulin, antibodies against Foxj1 (HPA005714, Sigma‒ Aldrich, 1:150) and NG2 (neuron-glial antigen 2, AB5320, Merck Millipore, Burlington, MA, USA, 1:400) were used.

The explants used to study the stem cell therapy effect with the Evans Blue assay were processed as described below.

### Bone marrow-derived mesenchymal stem cell isolation and culture

Bone marrow-derived MSCs were obtained from young male and female mice (20-24 days old) and characterized as previously described^43^. Briefly, bone marrow-derived MSCs were selected according to their plastic adhesion *in vitro*, their positive expression of CD44, CD73 and CD90, their negative expression of CD34 and CD45, and their trilineage differentiation capacity^27^. Dulbecco’s modified Eagle’s medium (DMEM, D5546, Sigma‒ Aldrich) containing 1% penicillin/streptomycin, 0.5% amphotericin B, 6.25% L-glutamine, and 10% fetal bovine serum (FBS, F7524, Sigma‒Aldrich) was used as the culture medium. After approximately 80% confluence at passage one, the cells were detached with trypsin/ethylenediaminetetraacetic acid (EDTA; T3924, Sigma‒Aldrich), centrifuged and resuspended in saline serum to be used to pretreat explants or primary cultures.

### Neural stem cell isolation and culture

The subventricular zone (SVZ) of the rostral-dorsal striatal wall in newborn male and female mice was dissected out and mechanically dissociated in DMEM/F-12 containing 100 units/mL penicillin‒streptomycin (P0781, Sigma‒Aldrich) to obtain NSCs, as described in other investigations^101, 115^. After centrifugation for 2 minutes at 600 g, the pellet was resuspended and placed overnight in uncoated plastic tissue-culture dishes in N5 medium constituted by DMEM/F-12 (Gibco 21331020, Thermo Fisher) containing N2 supplements (Gibco 17502001, Thermo Fisher), 35 μg/mL bovine pituitary extract (Gibco 13028014, Thermo Fisher), 5% FBS (F7524, Sigma‒Aldrich), 40 ng/mL EGF (epidermal growth factor, AF-100-15, PeproTech, Rocky Hill, New Jersey, USA) and 40 ng/mL bFGF (basic fibroblast growth factor, 130-093-564, Miltenyi Biotech, Bergisch Gladbach, Germany). New EGF and bFGF were added every 48 hours. For experimentation, cells were used at passage 2.

### Ependymal progenitor cell collection

EpPs were obtained from newborn male and female mice. In this case, the SVZ of the medial-dorsal striatal ventricle wall was dissected out and mechanically dissociated in DMEM-high glucose (glucose 4500 mg/L, D6546, Sigma‒Aldrich) with 10% FBS (F7524, Sigma-Aldrich), 1% L-glutamine (Gibco 25030081, Thermo Fisher), and 1% penicillin/streptomycin. Cells were plated at 500,000 cells/mL using the same media on poly-D-lysine (P6407, Sigma‒Aldrich)-coated round coverslips and incubated under standard cell culture conditions for 12 hours. The medium was switched entirely to fresh medium containing 2% FBS for another 12 hours. Astrocytes do not attach to plates in these latter conditions^101, 116^; thus, these cells were eliminated when the medium was switched. Therefore, in this case, only EpPs were attached to the plates.

### Stem cell transplantation in ventricular wall explants

Before application, NSCs, EpPs, and MSCs were labeled *in vitro* by adding green or red fluorescent cell tracker dyes (C2925, Thermo Fisher; and SCT107, Sigma‒Aldrich) following the manufactureŕs instructions.

Fluorescently labeled NSCs, EpPs and MSCs were gently detached from plates using trypsin/EDTA, centrifuged and resuspended in their respective media (NSCs and EpPs, 5,000 cells/µl; MSCs 5,000 cells/µl). Then, the stem cells were transplanted by accommodating them on the surface of the lateral ventricle explants using a Hamilton syringe (26-gauge, 1 µl/per explant) to avoid any damage to the surface of the explant and thus any additional alteration of the tissue.

Experiments were performed with 3 explants per experimental condition and replicated five times (n = 5), with a total of 15 explants per experimental condition.

For the analysis of the results, each explant was initially independently studied. Pictures were taken, and cells and markers were quantified. For each explant, a mean marker was obtained. For each replicate, we calculated the mean of the three explants. For the different statistical tests represented in the figures, the mean and standard deviation of the mean were calculated using the final value of each replica (n = 5).

### In vitro differentiation of ependymal progenitor cells under the effect of blood components

EpPs, obtained as described above, were plated at 500,000 cells/mL on poly-D-lysine-coated coverslips in 6-well culture plates (5 coverslips per well). Cells were incubated under standard cell culture conditions for 12 hours in high glucose DMEM with 10% FBS, 1% L-glutamine, and 1% penicillin/streptomycin. The medium was switched entirely to a new medium containing 2% FBS for another nine days. The medium was partially replaced every 48 hours.

Twenty-four hours after the medium was switched to 2% FBS, the EpP culture was treated and maintained with blood, blood serum, or TNFα. Blood and blood serum were obtained from newborn mice according to the above-described protocol and was applied to 500,000 cells/mL, and these treatments were applied once, 24 hours after the medium was switched to 2% FBS. TNFα (Genetex, Irvine, CA, USA) was applied daily at 50 ng/mL for a maximum of 9 days. In the different treatments, at 3, 6, and 9 days, the cells were fixed with 4% paraformaldehyde diluted in 0.1 M, pH 7.2 PB for 30 minutes at 4 °C and immunostained to detect Foxj1 protein and cilia (βIV-Tubulin) using unconjugated primary antibodies. Secondary fluorochrome-conjugated antibodies were used as described for brain sections. Nuclear staining was performed with DAPI. For negative controls, the primary antibodies were not added.

For analysis of the results, each well of the six-well plate was initially independently analyzed. Pictures were taken on each of the five coverslips, cells and markers were quantified, and an average was calculated per coverslip. Next, an average was calculated per well plate. Each condition was replicated 5 times. For the different statistical tests represented in the figures, the mean and standard deviation of the mean were calculated using the final value of each replicate (n = 5).

### In vitro differentiation recovery of ependymal progenitor cells in the presence of blood components and bone marrow-derived mesenchymal stem cells

EpPs, obtained as described above, were plated at 500,000 cells/mL on poly-D-lysine-coated coverslips in 6-well culture plates (5 coverslips per well). MSCs (20,000 cells/mL) were added once to the EpP primary culture 24 hours after the medium was switched to fresh medium containing 2% FBS and then maintained with the different treatments for nine days. Blood, blood serum, and the inflammatory cytokine TNFα were applied as described above, and cultures were processed, immunostained, and quantified as described in the *Image analysis and quantification* section.

For the analysis of the results, we proceeded as in the previous section. Each condition was replicated 7 times. For the different statistical tests, the mean and standard deviation of the mean represented in the figures were calculated using the final value of each replicate (n = 7).

### Evans Blue assay to test the brain parenchymal effect

Lateral ventricle wall explants from male and female mice with moderate GMH induced with collagenase were positioned on Millicell culture inserts in 6-well culture plates (3 explants/insert). After six hours *in vitro*, explants were transplanted with MSCs. Then, EpPs were transplanted 24 hours later. Transplantation was performed as described above. The medium was partially changed every 48 hours. After seven days of treatment, Evans Blue was applied as described in other studies^117, 118^. Then, tissue was fixed with 4% paraformaldehyde in 0.1 M PB, pH 7.2 at 4 °C for 24 hours.

Finally, explants were washed and analyzed under confocal microscopy at 650 nm based on the fluorescent properties of this compound. This colorant filled dead cells and edematous regions. Thus, the staining intensity indicated the grade of tissue damage and edematous status. Confocal images were transformed into greyscale, and every pixel obtained a value from 0 (black) to 255 (white). The highest intensity corresponds to a 255 value in the greyscale. The intensity of fluorescence in the grey scale was analyzed by ImageJ software. For each experimental condition, we used 6 explants.

### Cytokine content in culture media from treated explants

From the Evans Blue assay experiments described above, culture media were collected at the end of the experiment to detect cytokines after different treatments: cell culture media without explants as the baseline for the cytokine kit array (empty wells), nontreated explants (9 explants), MSC-pretreated explants (9 explants), and sequential transplantation of MSCs and EpPs in explants (9 explants). A mouse cytokine kit array (Abcam, Cambridge, UK; AB133995) was used to detect the expression of 62 cytokines according to the manufacturer’s instructions. Processed array membranes were scanned and analyzed using FIJI software 1.53q (NIH, USA). To obtain an accurate cytokine value for non-treated and treated explants, we subtracted the baseline expression of cytokines in the media with no explants for every case.

### Image analysis and quantification

Confocal images were acquired with SP8 and Stellaris 8 confocal laser microscopes (Leica, Wetzlar, Germany). For each experiment, immunofluorescence images were obtained in batches with control and experimental samples imaged under identical instrument settings. The ependyma-denuded ventricle surface percentage was quantified from micrographs (3-6 brain sections per animal) scanned under fluorescence in an Olympus VS120 microscope (Olympus, Tokyo, Japan) with a 10x objective.

Experiments were performed with 15 explants per experimental condition. Pictures were taken with a 20x objective in a Leica laser confocal SP8 microscope, and quantification was manually and blindly performed using the confocal images.

For the *in vitro* primary EpP culture, the experiments were performed with 7-9 wells per condition. In each well, 4-5 coverslips and 9-15 pictures were taken per coverslip, depending on the needed magnification for the experiment. Quantification was manually and blindly performed using confocal images.

### Statistics

Samples were analyzed using GraphPad 9.2.0 (GraphPad Software, San Diego, CA, USA) and Microsoft Excel 16.71. The required sample size was estimated according to standard deviations or significances. Normality was assessed with Shapiro–Wilk and D’Agostino’s K-squared tests. test was applied for hypothesis testing. For Student’s t test when the F probability from Student’s t test was < 0.05, the variance was considered unequal. For Student’s t test or the Wilcoxon-Mann Whitney test, two-tailed test was used. P < 0.05 based on both tests was considered statistically significant.

### Materials and Data Availability

The data that support the findings of this study are available from the corresponding author upon reasonable request.

